# Totipotency and high plasticity in an embryo with a stereotyped, invariant cleavage program

**DOI:** 10.1101/2025.02.12.637942

**Authors:** Amber Q. Rock, Mansi Srivastava

## Abstract

Animal embryos begin as totipotent zygotes, which undergo cell divisions and produce progeny with restricted fate potentials over time. However, the timing of when totipotency is lost and the processes through which embryonic cells acquire fates vary across species. Embryos with invariant cleavage programs, *e.g.* of nematodes and spiralians, tend to show early restriction of blastomere potency and limited robustness to perturbation, particularly after asymmetric cleavages have occurred. In contrast, embryos with variant cleavage programs, *e.g.* of vertebrates, tend to specify fates later in development and correspondingly show higher plasticity at early stages. Here, we investigate the embryos of the acoel *Hofstenia miamia*, which represents an understudied phylum (Xenacoelomorpha) that is distantly related to well-studied developmental systems. Given the invariant ‘duet’ cleavage program observed in *H. miamia* embryos, we found unexpected robustness in this species. Isolated 4-cell stage macromeres, the products of an asymmetric, fate specifying cleavage, were totipotent, forming whole organisms upon isolation. Notably, these isolated macromeres produced pharyngeal and neuronal tissues, which they do not produce during normal development. This assay is highly reproducible and can be done at high throughput in *H. miamia*, making this species an ideal system to investigate the causes of totipotency after specification. Photoconversion-based lineage tracing revealed that rescued cell types are not merely replaced by neoblasts, the adult pluripotent stem cells in *H. miamia*, suggesting that the macromere’s totipotency is the result of changes in the fate potentials of early embryonic cells. Remarkably, all blastomeres at the 8-cell stage were capable of reprogramming their fates in embryo reconstitution assays. By assembling different subsets of 8-cell stage blastomeres, none of which are totipotent on their own, we determined that a minimal unit of two blastomeres, one macromere that produces gut and neoblasts and one micromere that is specified to produce muscle and epidermis, was sufficient to develop into a hatchling worm. Future studies of this system could identify the precise mechanisms that can enable tremendous plasticity, including post-zygotic totipotency, in an embryo with well-defined cellular lineages.

## Introduction

The zygote ultimately produces and adult animal with many distinct cells and tissues, but the cellular processes through which differential fates are specified during embryogenesis can vary greatly across species. For example, in embryos with invariant cleavage programs, blastomeres show fixed fates at early stages of development (e.g. in nematodes, ascidians, spiralians) (Conklin, 1905; Jeffery, 2001; Laufer et al., 1980; Maduro, 2010; Nishida, 2005; Sulston et al., 1983). Conversely, in embryos with variant cleavage programs, fate specification occurs later in development, with early blastomeres taking on different fates embryo to embryo (e.g. in cephalochordates, vertebrates) (Benirschke, 2009; Holland & Holland, 2007; Katayama et al., 2010). Notably, these different modes of development broadly correspond to different levels of plasticity and/or robustness. Many embryos with invariant cleavage are determinate, i.e. they are unable to recover cell fates lost upon blastomere ablation or isolation, whereas most embryos with variant cleavage are indeterminate and can recover cell fates lost to perturbation. Some vertebrate embryos show post-zygotic totipotency, where isolated blastomeres develop into whole animals. (Bedzhov et al., 2014; Davidson, 1990, 1991; Lawrence & Levine, 2006; McCain & Cather, 1989). Here, we characterize embryonic plasticity in an acoel worm that deviates from this pattern, showing unexpectedly high potency in early embryonic cells despite having invariant cleavage and a stereotyped blastomere fate map.

As likely early-diverging bilaterians (Cannon et al., 2016; Edgecombe et al., 2011; Egger et al., 2009; Hejnol et al., 2009; Kapli et al., 2021; Kapli & Telford, 2020; Marlétaz et al., 2019; Philippe et al., 2007, 2011; Ruiz-Trillo et al., 2004), acoels (Phylum: Xenacoelomorpha) are distantly-related to spiralians, yet their embryonic cleavage program is broadly reminiscent of spiral cleavage (Figure 1A). In contrast to spiralian embryos, which have four quadrants of macromeres that serially produce smaller micromeres toward the animal pole, acoel embryos undergo “duet” cleavage, with two duets of macromeres that produce micromeres toward the animal pole (Boyer et al., 1996; J. Q. Henry et al., 2000; Kimura et al., 2021). Lineage tracing studies of two distantly-related acoels, *Neochildia fusca* and *Hofstenia miamia*, suggest that the invariant duet cleavage program results in a stereotyped fate map of early cleavage that is consistent across embryos, albeit with some differences between species (J. Q. Henry et al., 2000; Kimura et al., 2022). Experimental studies in the acoel *Childia groenlandica*, which displays a variation of duet cleavage, showed that embryos are able to complete development upon ablation of one of the two blastomeres at the 2-cell stage (Boyer, 1986). However, no other data investigating the plasticity and robustness of xenacoelomorph embryos have been reported, leaving a gap in knowledge for this major lineage of animals. The three-banded panther worm, *H. miamia*, is a lab-tractable acoel (Ricci & Srivastava, 2021; Srivastava, 2022; Srivastava et al., 2014) with a known blastomere fate map (Kimura et al., 2021, 2022) that produces plentiful embryos. Therefore, we sought to rigorously assess plasticity in this system, particularly after asymmetric, fate assigning cleavages have occurred.

**Figure 1:**
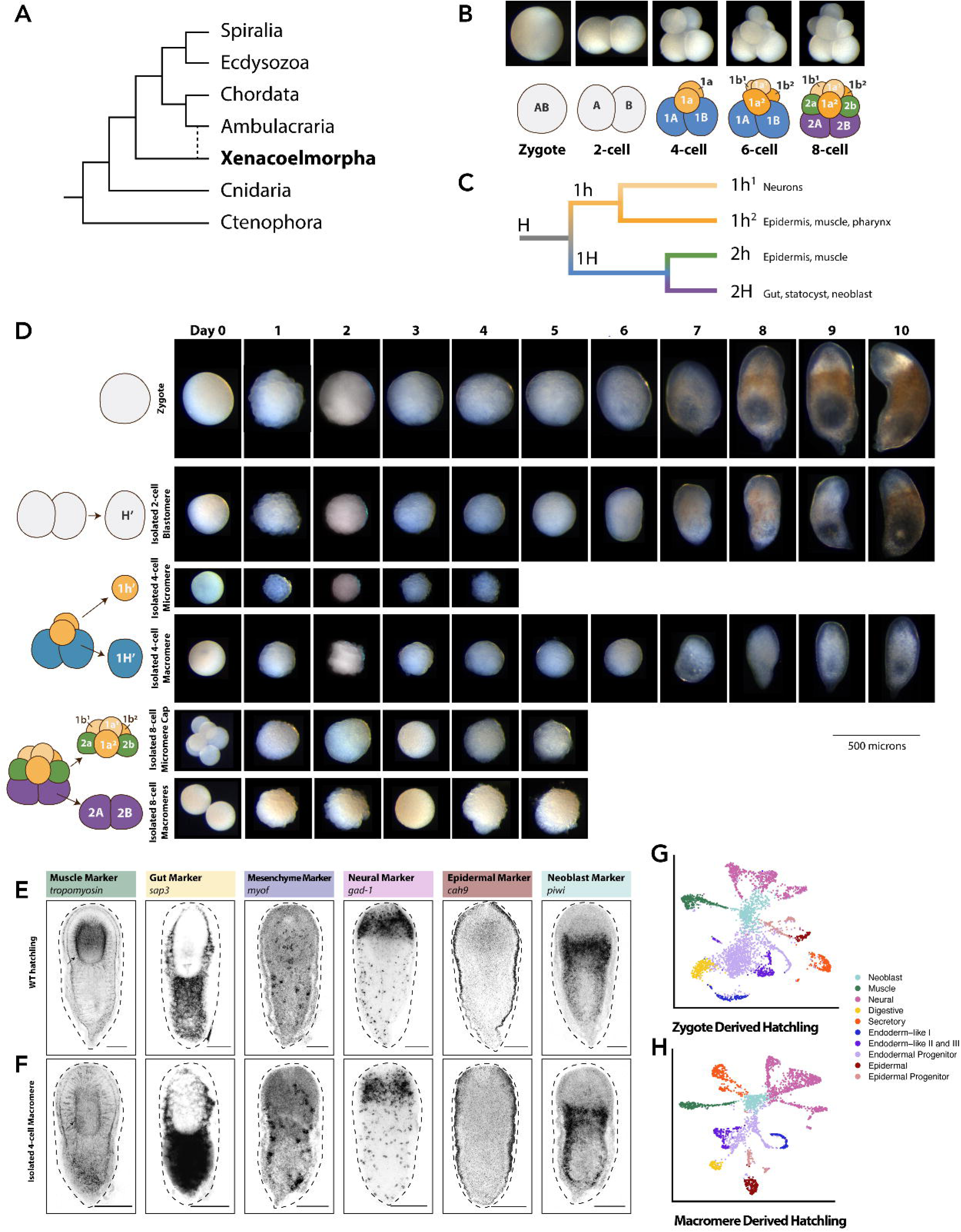
Isolated 4-cell macromeres develop into hatchling worms after a fate specifying cleavage. (A) Phylogenetic tree highlighting the position of Xenacoelomorpha. Dashed line indicates suggested alternative position. (B) Brightfield images and schematic diagram showing the stereotyped cleavage pattern of the dechorionated *H. miamia* embryo. The zygote undergoes its first cleavage, forming the equal sized blastomeres, A and B (2-cell stage). Each of those blastomeres next undergoes an asymmetric division, creating smaller cells at the animal pole called micromeres 1a and 1b (yellow), and larger cells at the vegetal pole called macromeres, 1A and 1B (4-cell stage). The first duet micromeres, 1a and 1b, divide, producing 1a^1^ and 1b^1^ at the animal pole and 1a^2^ and 1b^2^ vegetally. The macromeres, 1A and 1B (blue), then divide asymmetrically, creating a second set of micromeres, 2a and 2b (green), with the remaining macromere becoming 2A and 2B (purple) (8-cell stage). (C) Lineage tree of blastomeres beginning with the 2-cell stage blastomeres (A and B) notated as ‘H’ in reference to the A and B blastomeres corresponding to left and rights halves of the animal and being indistinguishable to the eye. At the 4-cell stage, the micromeres are the sole source of neuronal and pharyngeal tissue and contribute to muscle and epidermal tissue. The macromeres are the sole source of the gut, statocyst, and neoblasts, and contribute to muscle and epidermal tissue. At the 8-cell stage, the fates of the 4-cell stage macromere, 1H, are partitioned between 2h and 2H upon division, with 2h giving rise to epidermis and muscle, and 2H giving rise to gut, statocyst, and neoblasts. (D) Developmental time course of isolated blastomeres. From top to bottom, dechorionated zygote as control, isolated 2-cell blastomere, isolated 4-cell macromere (1H’), isolated 4-cell micromere (1a’), isolated 8-cell micromere cap (1h^1^, 1h^2^, 2h), and isolated 8-cell macromeres (2H). Scale bar 500 µm. Dechorionated zygotes develop into hatchling worms (n=313/397). Isolated 2-cell blastomeres develop into hatchling worms (n=221/263). Isolated 4-cell macromeres develop into hatchling worms (n=366/434). Isolated 4-cell micromeres fail to develop into hatchling worms (n=116/116). Individual isolated 8-cell macromeres fail to develop into hatchling worms (n = 68/68). Isolated 8-cell micromere cap fail to develop into hatchling worms (n=64/64). (E, F) Fluorescent *in situ hybridization* of major cell type markers in the hatchlings derived from dechorionated zygote controls (n=4/4, replicates per gene) and the hatchling derived 4-cell macromeres (n=5/5, replicates per gene) Black arrow denotes pharynx in *tropomyosin* FISH. (G, H) Uniform manifold approximation and projection (UMAP) plots representing scRNA-seq data from hatchlings derived from dechorionated zygote controls (n = 88) and hatchlings derived from isolated 4-cell macromeres (n=107). Cells colored by cell type.

Plasticity of blastomeres, *i.e.*, the ability of embryonic cells to adopt new fates upon perturbation, can change over time in the same embryo, with asymmetric cleavages often corresponding to reduction in plasticity. For example, the symmetric cleavages in sea urchin embryos produce blastomeres that are potent to produce whole animals, but subsequent asymmetric cleavages result in blastomeres that cannot compensate for the loss of cell fates upon isolation (Davidson et al., 1998; Livingston & Wilt, 1990). Strikingly, we found in *H. miamia* that at the 4-cell stage, macromeres, the larger progeny of the first asymmetric cleavage, are totipotent. We found that the recovery of neuronal and pharyngeal tissues, fates that are normally derived from the micromere lineage, is driven by transfating or reprogramming of early embryonic cells. At the 8-cell stage, no blastomere is totipotent or sufficient to complete development on its own, yet we found that all blastomeres were competent to adopt new fates in embryo reconstitution assays. This suggested that the *H. miamia* embryo relies on conditional specification to establish cell fate. Further, we determined the minimum set of blastomeres necessary for development at the 8-cell stage: the 8-cell stage macromere is necessary, and the addition of a micromere that is specified to make epidermis and muscle is sufficient for development. Altogether, these experiments reveal unexpected plasticity, including post-zygotic totipotency after an asymmetric cleavage, in an embryo with invariant cleavage.

## Results

### A macromere isolated from a 4-cell stage embryo is totipotent, and develops into an adult animal with all major cell types

Early *H. miamia* embryos undergo a stereotyped cleavage program with an invariant fate map (Figure 1B, C) (Kimura et al., 2021, 2022). Briefly, the zygote undergoes a symmetric cleavage, creating equally sized blastomeres, referred to as A and B, at the 2-cell stage. Each cell then undergoes an asymmetric division, yielding smaller micromeres (1a, 1b) toward the animal pole and larger macromeres (1A and 1B) toward the vegetal pole, reaching the 4-cell stage. Lineage tracing showed that the micromeres give rise to neurons and pharynx, and a majority of the epidermis and muscle. The macromeres give rise to the gut, statocyst, and neoblasts, the adult pluripotent stem cell population that facilitates regeneration and homeostasis in adult worm (Srivastava et al., 2014), as well as contributing to the epidermis and muscle. To reach the 8-cell stage, each micromere then produces progeny with increasingly restricted fates: smaller animal micromeres (1a^1^ and 1b^1^) give rise to the nervous system and larger vegetal micromeres (1a^2^ and 1b^2^) generate pharyngeal, epidermal, and muscle fates. Macromeres divide asymmetrically – forming new micromeres (2a and 2b) that also produce epidermal and muscle fates and new macromeres (2A and 2B) that make gut, statocyst, and neoblast fates. The A and B sides of the embryos are visually indistinguishable from each other, and undergo cleavage in the same sequence, producing cells with the same fate potentials. Therefore, similar to the use of the letter Q to refer to A, B, C, or D quartets in spiralian embryos (Guralnick & Lindberg, 2001; Lambert, 2010), we use the letter ‘H’, for halves, to refer to symmetrical blastomeres on the A and B sides (Supplemental Figure 1A). Further, we use the prime symbol to indicate blastomeres that have been isolated.

To assess the developmental potential of early embryonic blastomeres, we dechorionated embryos and 1) isolated individual micromeres and macromeres at the 4-cell stage (1h’ and 1H’) and, 2) separated the micromere cap (1a^1^, 1b^1^, 1a^2^, 1b^2^, 2a, and 2b) and the paired macromeres (2A and 2B) at the 8-cell stage, and cultured them to continue development until hatching or failure (Figure 1D). Dechorionated zygotes developed into hatchling worms, demonstrating that the chorion is not necessary for development. Additionally, isolated blastomeres at the 2-cell stage, H’, developed into hatchling worms, demonstrating that dissection does not inhibit development. At the 4-cell stage, which forms as a result of the first asymmetric and fate-assigning cleavage, we found that isolated single macromeres, 1H’, but not isolated micromeres, 1h’, developed into hatchling worms (Figure 1D). Survival rates were similar between the two halves of the embryo (A and B), suggesting each half is equipotent. Isolated 4-cell stage macromeres also showed similar hatching rates to dechorionated zygotes and isolated 2-cell blastomere halves (H’) (Supplemental Figure 1B). The 4-cell stage micromeres (1h’) did not complete development upon isolation, despite undergoing cleavage for 4-days post dissection. At the 8-cell stage, neither the micromere cap nor the paired macromeres were sufficient for complete development, although they both continued to cleave for 5 days.

To determine if hatchling worms derived from isolated blastomeres were comparable to hatchlings derived from zygotes, we performed fluorescent *in situ* hybridization (FISH) for markers of major cell types (Hulett et al., 2020; Srivastava et al., 2014) and found that all major cell types are present in the 1H’-derived hatchlings (Figure 1E-F, Supplemental Figure 1C). Additionally, we performed single-cell RNA sequencing (scRNA-seq) on 10-day old hatchlings derived from both dechorionated zygotes and 1H’ and found that both datasets contained all major cell types expected based on previous scRNA-seq data (Hulett et al., 2023) (Figure 2G-H, Supplemental Figure 1D). Notably, although 1H macromeres don’t contribute to neural and pharyngeal fates in unperturbed embryos, they were able to produce these cell types, as indicated by FISH and scRNA-seq. Further, with more time to feed, grow and mature, the adult worms derived from 1H’ laid embryos and exhibited normal regeneration (n=4/4), indicating they were bonafide worms. Altogether, these assays show that worms formed by isolated 1H’ macromeres were indistinguishable from wild type worms, confirming that the isolated 4-cell macromere shows post-zygotic totipotency. To our knowledge, this is the first report of a single blastomere produced by an asymmetric and fate-assigning cleavage exhibiting totipotency. Given that the isolated macromere 1H’ is totipotent, we sought to identify the source of pharyngeal and neuronal cell types, which are normally derived solely from micromeres (Rock & Srivastava, 2025).

**Figure 2:**
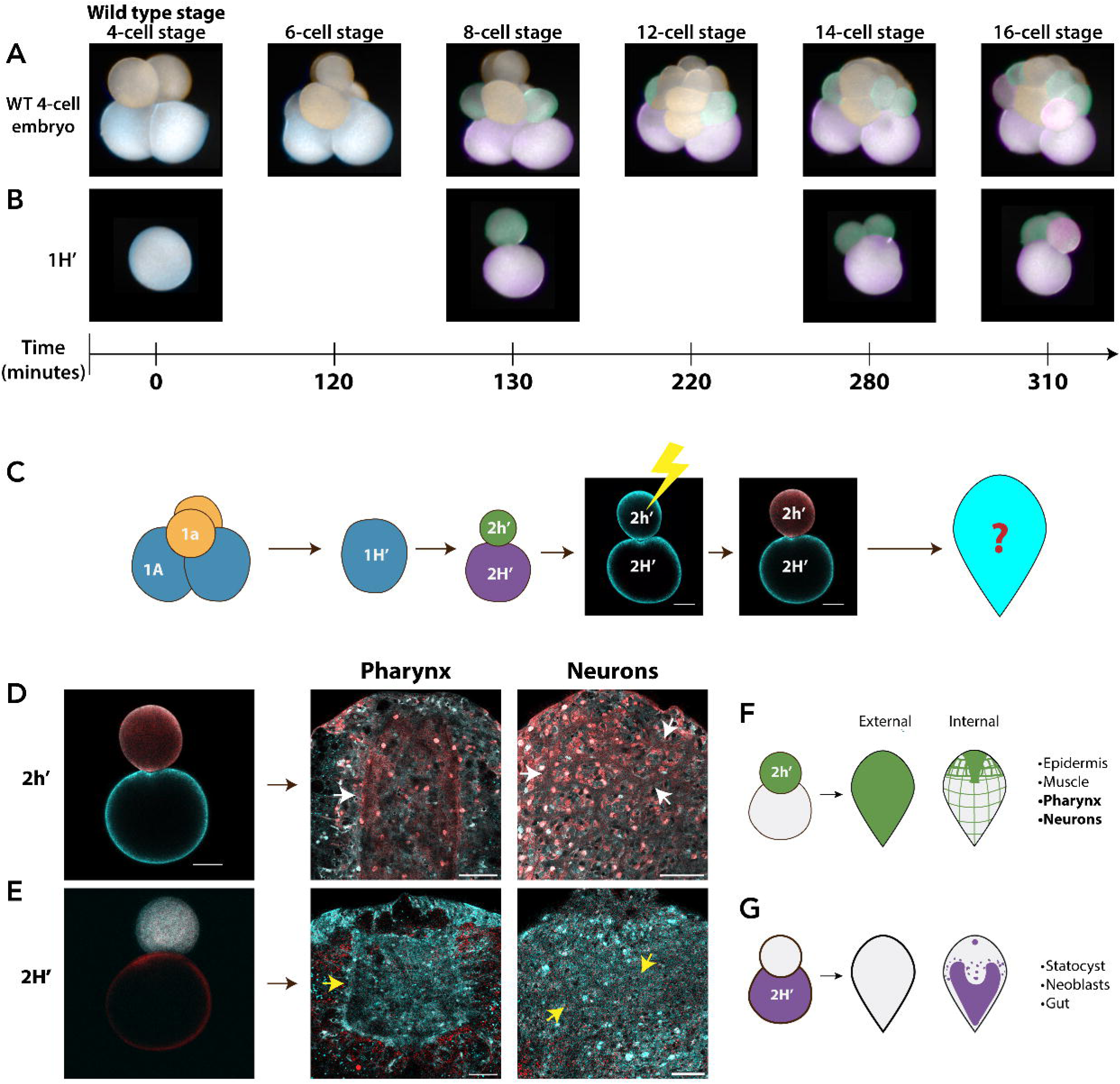
The isolated 4-cell stage macromere, 1H’, continues its cleavage program. The blastomere 2h’ is the source of missing cell types. (A-B) Images of the cleavages until the 16-cell stage in the wild type embryo and the isolated 4-cell macromere, 1H’, immediately after isolation. 1H’ maintains the WT invariant cleavage program in isolation. Blastomeres are false colored based on blastomere identity. (C) Schematic of example experimental design. 1H’ is isolated from a *tba1a*::Kaede embryo, allowed to cleave once, and blastomere 2h’ (or 2H’) is photoconverted. The embryo is allowed to develop until hatchling and then assessed for labeled cell types. (D) Results of lineage tracing show 1h’ gives rise to pharynx and neurons (n=6/6). Image duplicated from Figure 2C. (E) Resulting lineage tracing of 1H’ does not gives rise to pharynx or neurons (n=7/7). Photoconverted Kaede (red) labels pharyngeal, neural, muscle, and epidermal cells. (F, G) Schematic of the cell types formed by 2h’ and 2H’. 2h’ is the source of 1h^1^ and 1h^2^-specific fates in the isolated 4-cell macromere. Photoconverted Kaede (magenta) and unconverted Kaede (cyan). Arrows point to labeled (white) and unlabeled (yellow) cell types. Scale bar 50 microns.

### Expanded developmental potential of early blastomeres, not neoblasts, drives the development of missing cell types in isolated 4-cell stage macromere, 1H’

To determine the mode through which 1H’ recovers neural and pharyngeal fates, we first made timelapse movies to compare the cleavage pattern of 1H’ relative to 1H blastomeres that are in their native context in dechorionated zygotes. We found that the isolated 1H’ cell continues to produce progeny with the same timing, size, and order as unperturbed 1H cells (Figure 2A-B, Supplemental Figure 2A-B). This suggested that the blastomere does not respond to isolation by altering its cleavage program and may continue to produce progeny with similar identities to those in unperturbed contexts.

Given that in unperturbed embryos 1H blastomeres give rise to neoblasts (Kimura et al., 2022), we sought to determine if neoblasts were the source of the cell types that are normally made by the 1h micromeres (neurons and pharynx). We employed lineage tracing using a transgenic line (Tuba::Kaede) that produces embryos with all cells expressing the photoconvertible protein Kaede (Kimura et al., 2022). We first isolated blastomeres from 4-cell stage Kaede^+^ embryos (1H’) and allowed them to undergo one cleavage. We then photoconverted either the micromere, 2h’, or the macromere, 2H’, to label the cell and allowed the embryo to develop until hatchling, where it was then fixed and assessed for the labeling of different cell types (Figure 2C). We reasoned that if neoblasts are a source of neurons and pharynx, these tissues should be labeled red in hatchlings made by embryos with photoconverted 2H’ macromeres, as this cell produces the 3h’ micromere that resembles the neoblast progenitors 3a and 3b in wild type embryos (Supplemental Figure 2A-B). Strikingly, 2H’ did not contribute to neurons and pharyngeal cells. Instead, we found that 2h’ gives rise to pharyngeal, neuronal, epidermal, and muscle cell types (Figure 2D, E, Supplemental Figure C, D). Given that 2h’ is the equivalent of 2a and 2b, which in unperturbed embryos do not give rise to neurons or pharynx, our results suggest that isolation of 1H’ results in considerable expansion of the cell fates of the 2h’ micromeres (Figure 2F, G). Thus, the totipotency of 1H’ appears to be driven by a process akin to reprogramming, and not likely via a neoblast-based mode of replacement.

Our data show that despite the immediate maintenance of the same cleavage program after isolation, blastomere 1H’ expanded its developmental potential to create all cell types, including pharynx and neurons. We wondered if this property was unique to the macromeres of the 4-cell stage embryos. Therefore, to determine if the other blastomeres (4-cell stage micromeres, 8-cell stage micromere cap, and 8-cell stage paired macromeres) also showed signs of expanded fate potential despite being incapable of completing development, we isolated them and assessed their developmental potential to determine if their fates had been specified or if they were capable of developing cell types beyond the tissues they contribute to in unperturbed embryos.

### Micromere fate is specified upon cleavage whereas macromere fate appears to be specified later in development

We isolated single micromeres and macromeres at the 4-cell stage and then performed scRNA-sequencing on the resulting embryos at 2- and 3-days post dissection to determine if they could make any cell types outside of their original fates during development (Figure 3A). We created an updated version of the *H. miamia* transcriptome to enrich for transcripts that may be specific to developmental timepoints. The resulting scRNA-seq data, as well as wild type developmental time course scRNA-seq data (Kimura et al., 2022) were mapped to the new transcriptome and then integrated for downstream analysis. A uniform manifold approximation and projection (UMAP) integrating data from 2- and 3-days post lay (dpl) embryos from wild-type zygotes (Kimura et al., 2022), isolated micromeres, and isolated macromeres showed distinct occupancy of the UMAP space between micromere- and macromere-derived samples at 2-dpl. At 3dpl, macromere-derived embryos showed expanded occupancy of clusters, while micromere-derived embryos appeared restricted to the neural progenitor cluster (Figure 3B, C). Cell clusters were assigned identities based on expression of markers that were specific to clusters in wild type embryos at 2- and 3-dpl, as well as co-expression of known differentiated cell type markers. We subsetted each experimental condition and quantified the proportion of cells that were present in each cell type cluster (Figure 3C, D). We found that there were significant differences in cell type composition of micromere- and macromere-derived embryos relative to the wild type embryos (Chi-squared test of independence, p-value < 2.2e-16, Supplemental Figure 2A). Despite differences in the proportion of cell types, macromere-derived data showed the presence of all major cell identities by 3dpl, consistent with the ability to complete development. In contrast, micromere-derived data did not show major contributions to cell type clusters that micromeres do not contribute to in the wild type embryo, such as gut and secretory. This suggested that micromere fate becomes specified upon their birth at the 4-cell stage.

**Figure 3:**
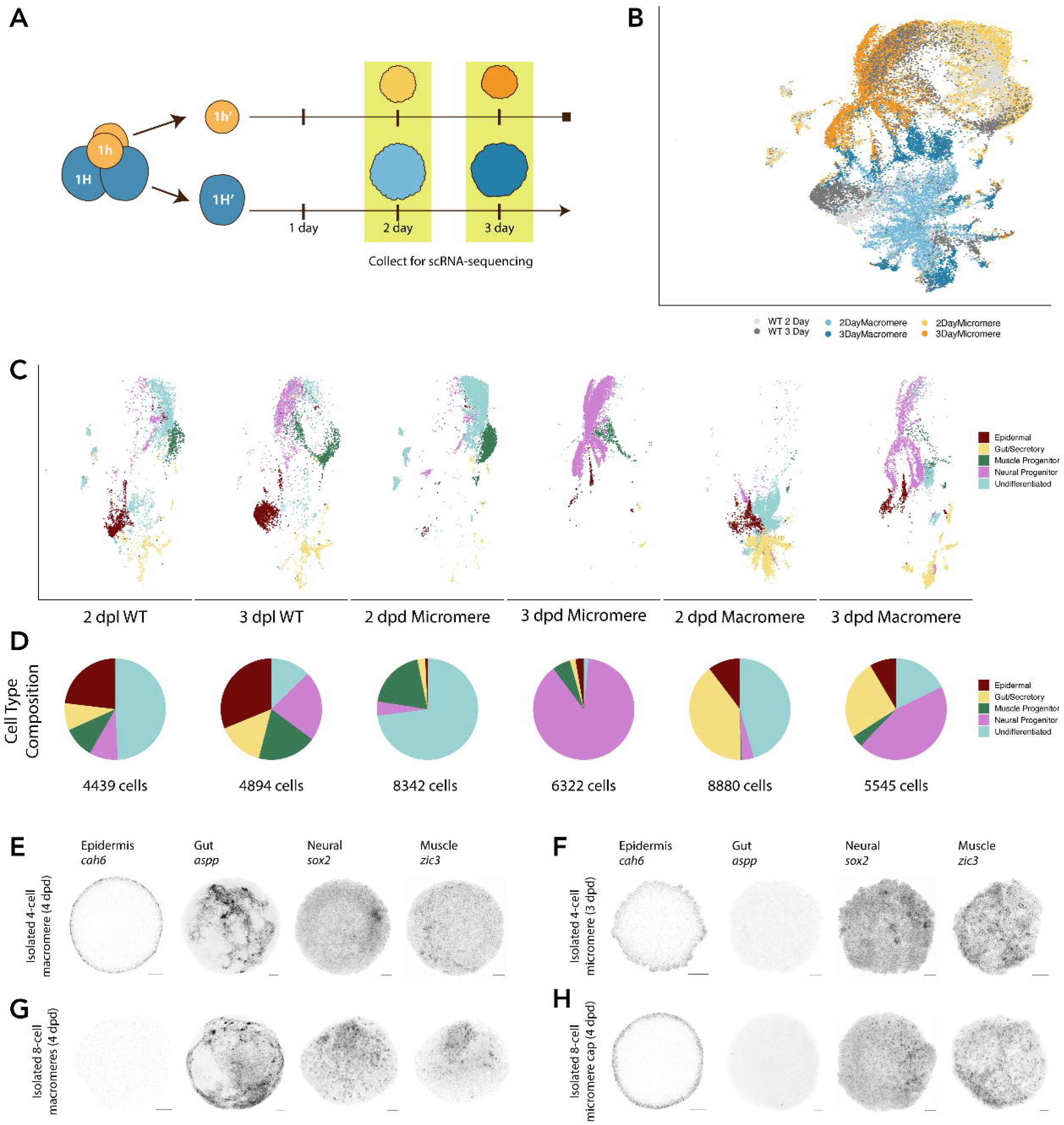
Micromeres are specified upon cleavage at the 4- and 8-cell stage while macromeres show expanded developmental potential in isolation. (A) Schematic of experimental design. 4-cell stage micromeres and macromeres were isolated and reared for 2 and 3 days. The embryos were then dissociated for scRNA-sequencing. (B) UMAP for an integrated scRNA-seq object with wild type 2- and 3-day post lay (dpl) embryos and 2- and 3-days post dissection (dpd) 4-cell micromeres and macromeres. Wild type embryo cells are colored in grays, micromere derived cells are colored in yellows and macromere derived cells are colored in blues. (C) Subset UMAPs for each experimental group with cells colored by cluster cell type identity (burgundy for epidermal, yellow for gut/secretory, green for muscle, pink for neural, and light blue for undifferentiated). (D) Pie charts depicting the proportion of cells in each cell type for each experimental group. The total number of cells for each group written is below pie chart. (E-H) FISH of cell type markers for epidermis, gut, neurons, and muscle in embryos derived from 4-dpd 4-cell macromeres, 8-cell macromeres, and 8-cell micromere caps, and 3-dpl 4-cell micromeres. Scale bar 50 microns. (E) Embryos from 4-cell macromeres express markers for all cell types (5 replicates per gene). (F, H) Embryos from 4- and 8-cell micromeres only express previously specified cell types (epidermis, neurons, and muscle) and not gut (6 and 4 replicates per gene, respectively). (G) Embryos from 8-cell macromeres give rise to gut, as well as neurons and muscle, which the cells do not contribute to in the wild type context, and do not give rise to epidermis (4 replicates per gene).

We then assessed the presence of both known (Bump et al., 2023; Kimura et al., 2021) and newly identified cell type markers via FISH in embryos derived from 4-cell stage micromeres and macromeres, (Figure 3E-F, Supplemental Figure 3D-E). We also assessed embryos derived from the 8-cell stage micromere caps and paired macromeres, as neither is sufficient for development (Figure 3G-H, Supplemental Figure 3F-G). We found that the totipotent 4-cell macromere produced an embryo that showed expression for all assayed cell types (Figure 3E, Supplemental Figure 3B-C, E). Both the 4-cell stage micromere and 8-cell stage micromere cap produced embryos that showed expression of markers for cell types it contributed to in the wild type context (epidermis, neurons, and muscle) cell types, but did not show expression for gut, demonstrating an inability to produce cell types outside of its assigned fate (Figure 3F, H, Supplemental Figure 3D, F). The embryos derived from the 8-cell stage macromeres showed expression of presumptive gut markers, a cell type it contributes to in the wild type embryo, as well as neural and muscle markers, cell types it does not normally give rise to, demonstrating an expansion of developmental potential of the cell in isolation (Figure 3G, Supplemental Figure G). This confirmed that micromeres are specified upon cleavage, whereas macromeres retain higher potency.

### All blastomeres at the 8-cell stage are plastic, showing expanded developmental capacity in embryo reconstitution assays

Although micromere fate is specified upon cleavage beginning at the 4-cell stage, it is possible that this specification is mutable in response to a perturbation, such as blastomere position. To determine the potency of blastomeres, we implemented an embryo reconstitution assay (Figure 4A): 1) The blastomeres of both an unlabeled, wild type embryo and a fluorescently labeled embryo, both at the 8-cell stage, where no blastomere is totipotent on its own, are carefully separated such that their identities can be tracked. 2) one pair of the fluorescently labeled blastomeres (either 1h^1^, 1h^2^, 2h, or 2H) is used to replace the corresponding blastomeres of the wild type embryo, such that two fluorescent ‘donor’ blastomeres and six ‘recipient’ unlabeled blastomeres together compose the full set of blastomeres normally present at the 8-cell stage. 3) These eight blastomeres are allowed to mix, settle and reaggregate at the bottom of a PCR tube. Presumably, the normal positions and cell contacts that would be found in a wild type embryo would be perturbed in these reconstituted embryos. 4) When the embryos develop into hatchlings, the resulting animals are fixed and assessed for fluorescently labeled cell types.

**Figure 4:**
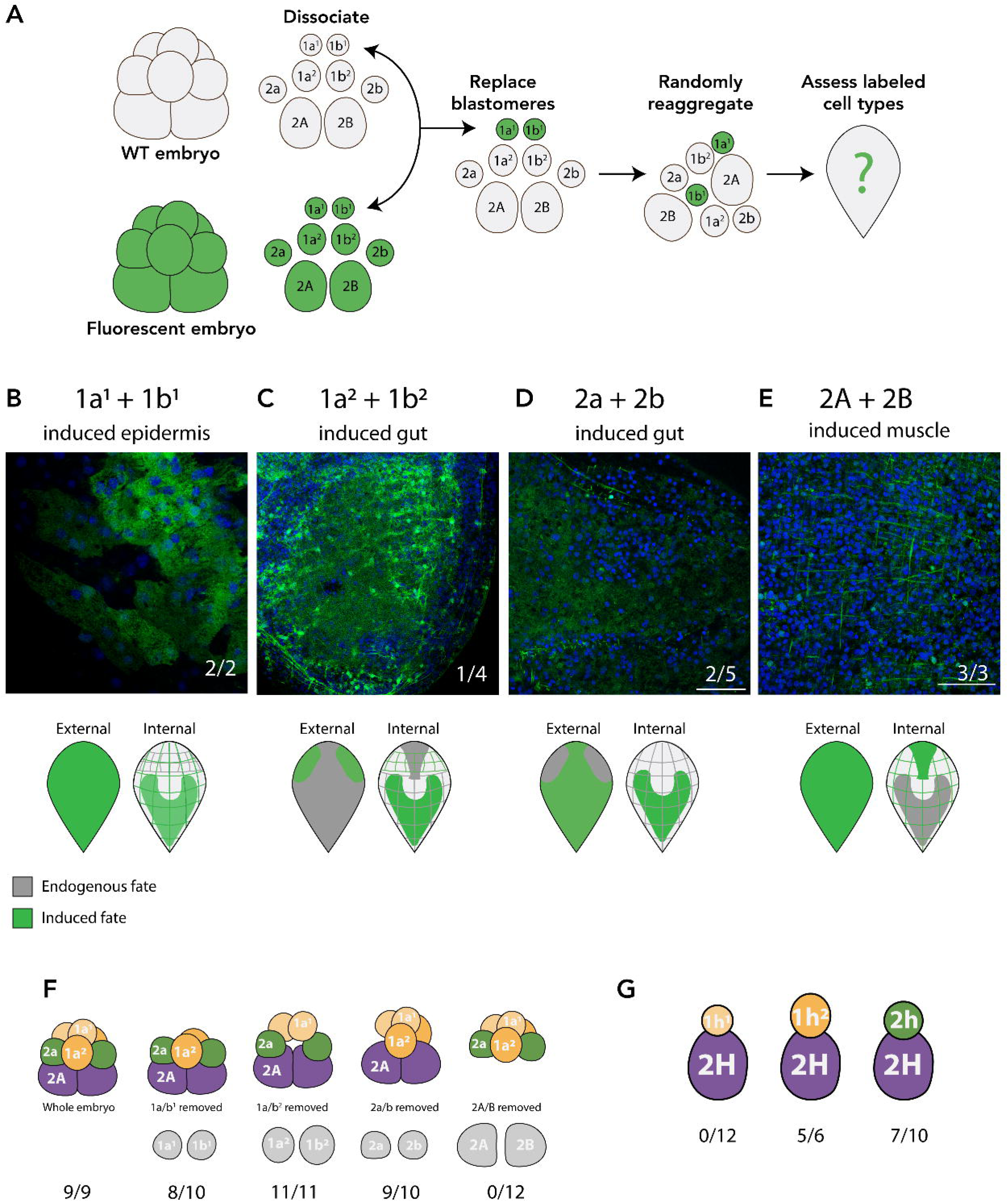
Blastomeres show expanded developmental potential when reaggregated. (A) Schematic diagram of the experimental design. A wild type embryo and fluorescently labeled embryo were both dissected. A pair of blastomeres from the wild type were removed and replaced by the labeled blastomeres. The eight blastomeres were randomly reaggregated and allowed to develop into a hatchling worm, which was the assessed for fluorescently labeled cell types. (B-E) Representative images of labeled cell types (green) and nuclei (blue) with schematics highlighting newly induced cell types (bottom). The number of replicates is indicated in the figure. In this perturbed context, all blastomeres give rise to cell types beyond their fate in the wild type embryo. (F) Schematic of experiments to determine necessary blastomeres at the 8-cell stage. Only the macromeres are necessary at the 8-cell stage. (G) Schematic of the minimum blastomeres necessary to produce a hatchling worm. A single 8-cell macromere (2H) and either a 1h^2^ or 2h micromere is sufficient to produce a hatchling worm (n= 5/6 and 7/10, respectively).

We found that all blastomeres showed an expansion of the cell types they contribute to relative to their endogenous fates in unperturbed embryos (Figure 4B-E, Supplemental Figure 4 A-D). The 8-cell stage macromeres, 2A and 2B, which normally form the gut, statocyst, and neoblasts in the wild type embryo give rise to epidermal, muscle, and pharyngeal tissue in the reconstitution assay. This is perhaps not surprising, as these cells do show expanded fates upon isolation, developing muscle and neurons, though they do not give rise to either cell type in the 1H’ context (Figure 2E). It is of note that micromeres 1a^1^ + 1b^1^, 1a^2^ + 1b^2^, or 2a + 2b, which have already undergone specification and cannot make any other cell type in isolation, upon reconstitution are able to contribute to fates they do not normally produce, such as gut. For example, 1a^1^ + 1b^1^ normally only give rise to neurons, but contribute to epidermis, muscle, and gut when reconstituted (Supplemental Figure 4A). This suggests that these blastomeres are competent and responsive to signaling, demonstrating the blastomeres’ plasticity and the reversibility of the cell’s specification. These experiments show that although no blastomere is pluripotent at the 8-cell stage, all blastomeres remain plastic after specification and show expanded developmental potential in response to reaggregation.

### The 8-cell stage macromere is necessary, but insufficient for development, and development can be rescued with addition of some 8-cell stage micromeres

Although many blastomeres proved capable of contributing to all major cell types in the reconstitution assay, no single 8-cell blastomere could develop into a complete worm when isolated. This made us wonder whether any set of blastomeres were necessary and/or sufficient for complete development, with the objective of defining the minimal components necessary for embryonic development in *H. miamia*.

To determine if any 8-cell stage blastomeres were necessary for development, we removed each pair of blastomeres in an embryo and allowed the embryo comprising the remaining six blastomeres to develop until failure (Figure 4F). When any micromere pair (1a^1^ + 1b^1^, 1a^2^ + 1b^2^, or 2a + 2b) was removed, the remaining embryo developed into a worm. When the macromeres (2A + 2B) were removed, the embryo failed to develop. Therefore, we conclude that only the macromeres are necessary for complete development. This is consistent with the specification of micromeres; without the macromeres they cannot develop necessary exogenous cell types, such as gut.

Given that the 8-cell stage macromeres are necessary but insufficient in isolation for development, we sought to determine which micromeres would be sufficient to enable complete development. Because 1H’ is totipotent and later is composed of the blastomeres 2H and 2h, we tested whether 2H and another micromere (1h^1^, 1h^2^, or 2h) would be sufficient for the complete development of the embryo (Figure 4G). Interestingly, we found that both 2H + 2h and 2H + 1h^2^, but not 2H + 1h^1^, could develop into a hatchling worm. This could be due to a necessity for the formation of an epidermis, as 1h^2^ and 2h both give rise to epidermis and muscle but 2H cannot create epidermis in isolation (Figure 3G). Alternatively, only 1h^2^ and 2h have endogenous contact with 2H during development, raising the possibility that there is something inherent to their cell-cell contact that is necessary for development. It is also possible that there is a signaling molecule found within these micromeres that is not present in 1h^1^.

## Discussion

Here we present a characterization of embryonic plasticity in *Hofstenia miamia*, an acoel worm whose embryos follow an invariant, stereotyped cleavage program. We found that the isolated 4-cell stage macromere, which is produced by an asymmetric cleavage where the fate of the sister micromere cell is specified, is totipotent. Despite never contributing to epidermal or pharyngeal lineages in the wild type embryo, this cell is able produce these tissues and ultimately makes a fully functional worm. This post-zygotic totipotency is distinct from adult, neoblast driven regeneration, as we found evidence for transfating or reprogramming of early blastomeres. Additionally, we also find high levels of cellular plasticity in the isolated 8-cell macromeres.

Post-zygotic totipotency has not yet been documented in an embryo after specification. For example, the crustacean *Parhyale hawaiensis* has an invariant cleavage program where fate appears to be autonomously specified when blastomeres are isolated and deleted blastomeres can only be compensated for from sister blastomeres that give rise to the same germ layer (Extavour, 2005; Gerberding et al., 2002; Price et al., 2010). Regulative capacity following the deletion of blastomeres is quite uncommon within protostomes, with the embryo from the tardigrade *Thulinia stephaniae* being a rare example, although that embryo undergoes variant cleavage without clear fate specification (Hejnol & Schnabel, 2005). Looking within deuterostomes, we see higher capacity to recover from blastomere deletion. For example, the sea urchin embryo undergoes two equal cleavages, creating four equally sized blastomeres. Each of these blastomeres, when isolated, can develop a complete, miniature embryo (Hörstadius, 1957). However, once the embryo undergoes an asymmetric cleavage during the fourth division, creating a tier of mesomeres, macromeres, and micromeres, each with their own fate specification, embryonic plasticity is substantially diminished. Neither the animal nor vegetal tier at this stage, nor the individual blastomere tiers is capable of completing development, with few instances of the cells being able to create cell types outside of what has been specified at that point (J. J. Henry et al., 1989; Livingston & Wilt, 1990; Wilt, 1987; Yamaguchi et al., 1994). These examples demonstrate the unique ability of the *H. miamia* embryo to maintain totipotency after a lineage specifying cleavage.

Our observation of unexpected plasticity in the *H. miamia* embryo raises the question of which mechanisms could maintain flexibility of fates in the context of a seemingly constrained fate map. Based on the embryo reconstitution experiment, we infer that cell-cell signaling is an important component of specification and differentiation of cell types in the *H. miamia* embryo. All embryos use a combination of both maternal determinants and conditional signaling to specify fates. Even in the highly autonomous ascidian embryo, signaling factors play an important role in differentiation (Kobayashi et al., 2003; Nishida & Sawada, 2001). There is also signaling in *C. elegans* embryos (Priess & Thomson, 1987). We suggest that cell contacts and communication have a large influence on the fates the *H. miamia* blastomeres take on. For example, the blastomere 2h normally only gives rise to epidermal and muscle fates in the wild type embryo. However, in the isolated 4-cell macromere 1H’, the resulting 2h blastomere now gives rise to neuronal and pharyngeal fates. Furthermore, when 2h is dissociated and reaggregated, it is capable of contributing to gut fates, demonstrating immense plasticity in differing conditions.

By investigating blastomeres’ necessity and sufficiency for embryonic development, we can begin to parse which aspects of early embryogenesis and specification are driven by inherent determinants or responsive signaling. At the 4- and 8-cell stage, macromeres are required for development, although only the 4-cell macromere is sufficient for development. We have determined that the minimum components necessary for development are an 8-cell macromere (2H) and either the 2h or 1h^2^ micromere, despite none of these blastomeres giving rise to neuronal lineages in the wild type embryo. Both 2h or 1h^2^ contribute to epidermis and muscle in the wild type embryo, a rare redundancy in fate in the *H. miamia* embryo. Although 2H can make muscle in isolation and muscle and epidermis when perturbed, it is possible that 2H alone cannot compensate for the missing cell types quickly enough, especially given that epidermis is one of the earliest cell types to differentiate (Kimura et al., 2021). It is possible that because neurons differentiate much later in development, the embryo has more time to compensate for this lineage’s loss.

Our study has revealed that the isolated 4-cell macromere, 1H’, is totipotent. This is a surprising example of post-zygotic totipotency after a fate specifying cleavage. Given the asymmetric divisions, there may be selective partitioning of maternal determinants that cause specification or allow plasticity in the blastomere. Future studies may reveal the maternal determinants and determine if there is conservation of mechanisms for plasticity in other animal embryos.

## ACKNOWLEDGMENTS

We thank Cassandra Extavour, Gonzalo Giribet, Kristen Koenig, Deirdre Lyons, Mark Martindale, Kara McKinley, Elaine Seaver, David Weisblat, Athula Wikramanayake, Jessica Whited, and members of the Srivastava Lab for insightful discussions. We thank Catriona Breen, Paul Bump, Vikram Chandra, and Allison Kann for their close reading of the manuscript. Amber Rock was supported by the Department of Organismic and Evolutionary Biology, the NSF-Simons Center for Mathematical and Statistical Analysis of Biology at Harvard, and the Harvard Quantitative Biology Initiative (#1764269). Mansi Srivastava was supported by the Smith Family Foundation Odyssey Award and NIH grant R35GM153252.

## METHODS

### Animal husbandry and embryo collection

Reproductive adult *H. miamia* worms were kept in plastic boxes containing approximately 1 liter of artificial sea water (37 ppt, pH 7.8-8.2, Instant Ocean Sea Salt) at 21°C. Worms were fed with brine shrimp *Artemia* sp. twice a week. Three days after being fed, the sea water was replaced, and detritus was wiped from the plastic boxes. All worms are progeny of random mating pairs from 120 sexually reproducing worms collected from Bermuda in 2010. Spontaneously laid embryos were collected with a glass pipette and then staged for experimental use.

### Embryo dechorionation and dissections

The chorion was removed from early-stage embryos to perform dissections. Embryos were agitated on an orbital shaker for 4-8 minutes while in a dechorionation solution (10mg/mL sodium thioglycolate, 0.1mg/mL pronase, and 10N NaOH in artificial seawater). Embryos that were fully dechorionated and intact were transferred to a dish coated with 1% agarose in artificial sea water. Blastomeres were isolated and removed using an eyelash tool. Single embryos/blastomeres were placed in individual wells of a 96-well plate in artificial sea water with 1:2000 penstrep. The water was replaced every three days, being careful not to fully remove all water, as cells will lyse if they touch the surface of the water. All plastic plates and pipettes were coated with 4% bovine serum albumin in artificial sea water to prevent sticking.

### Lineage tracing

Macromeres were isolated from a dechorionated 4-cell staged Tubulin:Kaede embryo. The blastomere was placed in a 4% bovine serum albumin coated ibidi µ-Slide (Cat. # 81506, ibidi) and given time to undergo a single division before an individual blastomere was photoconverted using a 405nm laser on a Leica SP8 confocal microscope, as previously reported (Ricci & Srivastava, 2021). The embryos were then transferred to a BSA coated 24-well plate with artificial sea water with 1:2000 penicillin and streptavidin antibiotics (Cat. # 15140122, Thermofisher Scientific). The water was replaced every three days until the embryos developed to hatchling stage, about 9 days. The resulting hatchling worms were then fixed with 4% paraformaldehyde in PBS overnight at 4 deg. Hatchlings were stained with DAPI then imaged on a Leica SP8 confocal microscope at 10x and 20x magnification.

### Blastomere replacement, dissociation, and embryo reconstitution

Wild type and *tba1a*::Kaede or *tba1a*::RFP embryos were dechorionated to serve as “donor” blastomeres (Ricci & Srivastava, 2021). The blastomeres were isolated while keeping trach of identity. Pairs of blastomeres (1a/b1, 1a/b2, 2a/b, and 2A/B) were replaced from the wild type embryo with Kaede or RFP blastomeres. The embryos were reaggregated in a 4% BSA coated PCR tube with artificial sea water and 1:2000 penstrep. Two days post reaggregation, the embryos were transferred to a BSA coated 24-well plate with artificial sea water with 1:2000 penstrep. The water was replaced every three days until the embryos developed to hatchling stage, about 9 days post-reaggregation. The resulting hatchling worms were then fixed with 4% paraformaldehyde in PBS overnight at 4 deg. Hatchlings were stained with DAPI then imaged on a Leica SP8 confocal microscope at 10x and 20x magnification.

### Probe synthesis and fluorescent *in situ* hybridization

Probe synthesis and fluorescent *in situ* hybridization were performed as previously described (Srivastava et al., 2014). Primer sequences for screened genes are provided in Supplementary Table 1. Embryos were dechorionated and dissected, if necessary, as described above and grown in pen-strep water until the appropriate stage. The following modifications were made to the protocol: Embryos were fixed in 4% PFA in PBST overnight at 4°C. All washed were done in 800 μL of solution. Embryos were permeabilized with 10 μg/ml Proteinase K and 0.1% SDS in PBST for 1 minute. The hybe solution was slightly modified containing: 50% Formamide (Invitrogen AM9342), 5X SSC (Corning 46-020-CM), 5% Dextran sulfate (Sigma D8906-100G), 500 μg/ml Torula RNA (Sigma R6625-25G), 1X Denhardt’s (Invitrogen 750018), 5 mM EDTA (Invitrogen AM9260G), 1% Tween-20 (Sigma P9416-100ML), and 50 μg/ml Heparin (Sigma H3393-50KU). The prehybe solution is the same as the hybe except the Dextran sulfate, Denhardt’s, and Heparin were replaced with water. Embryos counterstained with 1:2000 TO-PRO-3 Iodide (Cat. # T3605, Invitrogen), then washed with PBST before being imaged with a Leica SP8 confocal microscope.

### Transcriptome assembly

A *de novo* transcriptome was assembled using the pipeline developed by (Cunha & Giribet, 2019). New transcriptome assemblies were made from bulk RNA-seq data from early embryos (1, 2, 4, & 8 cell embryos) and from embryonic developmental stages (BioProject ascension code PRJNA603318) (Kimura et al., 2021) using three different transcriptome assembly programs (Trinity (both de novo and reference-based), StringTie, and Spades) (Bushmanova et al., 2019; Grabherr et al., 2011; Haas et al., 2013; Pertea et al., 2015). All assemblies were merged into a single assembly and ran through EviGene, which removes perfectly redundant genes and noncoding genes (Gilbert, 2013). The resulting genes were run through Transdecoder, which removed any genes that did not produce at least 100 aa product. Genes were then reverse complemented, when appropriate. The resulting genes were then mapped against the original bulk RNA datasets, individually. Any gene that had at least 5 TPM with any of the samples passed the filter. Those genes were then merged with the previous *H. miamia* transcriptome. CD-HIT was run and removed any genes that were 98%+ similar, retaining the longer of the two transcripts. The resulting transcripts were named using blastP to HumanSwissProt (<evalue 1e-3) (The UniProt Consortium, 2023). Genes that were newly assembled or had updated have a gene ID that begins with 982XXXXX. Transcriptome can be accessed online (NCBI Bioproject PRJNA1208142).

### inDrops single cell RNA collection and sequencing

Hatchlings developed from dechorionated zygotes (n=88), isolated 2-cell blastomeres (n=132), and isolated 4-cell macromeres (n=107) and 2-(n=180) and 3-(n=171) days post dissection 4-cell macromeres and 2-(n=168) and 3-(n=172) days post dissection 4-cell micromeres were dissociated and prepared for single cell inDrops encapsulation as previously described (Hulett et al., 2023). Dissociated cells were encapsulated using a microfluidics chip to minimize time in PBS before barcoding (Zilionis et al., 2017). Libraries were prepared by the Harvard University Single Cell Core and sequenced on an Illumina NextSeq500 using inDrops v3 design (75bp, paired).

### Single-cell processing and analysis

Reads were processed using the following github repositories: https://github.com/indrops/indrops and https://github.com/brianjohnhaas/indrops (which required python, bcl2fastq, RSEM, bowtie, samtools, pysam) and analyzed using Seurat v4.3.0.1. Outliers were filtered out based on low UMI counts. Cell types were assigned to clusters based on known adult cell type markers in the wild type 2- and 3-dpl datasets (Hulett et al., 2023). Clusters were labeled with the appropriate cell type prior to integration. Experimental embryonic time points were integrated with wild type embryonic time points (PRJNA888438) using Seurat. Chi-squared test of independence was used to determine differences in proportion of cell types found in the dataset for each experimental group. Early cell type markers used for *in situ* were selected based on wild type developmental single cell time course (Kimura et al., 2022). We identified genes that were exclusive markers to clusters in early developmental timepoints and co-expressed with a known cell type marker in at least one later point. Early cell type markers were further validated by examining the expression pattern via FISH in hatchling animals, where cell types are more easily distinguished by the shape and location of expression pattern.

## Data availability

Raw reads have been deposited to NCBI BioProject for early embryo bulk RNA-sequencing (PRJNA1208115) and inDrops single-cell sequencing from embryos and hatchlings (PRJNA1208470). The updated transcriptome has been deposited to NCBI BioProject (PRJNA1208142).

**Supplemental Figure 1.** (A) Survival curve from a subset dissected embryos. 75% of embryos from dechorionated zygotes (n = 48), 83% from isolated 2-cell blastomeres (n = 42), and 78% from isolated 4-cell macromeres (n = 61) develop into hatchling worms. All embryos derived from isolated 4-cell stage micromeres (n = 54), 8-cell stage micromere cap (n = 29), and 8-cell stage macromeres (n=28) failed to complete development and died. (B) FISH of major cell type markers in hatchlings derived from isolated 2-cell blastomeres. (C) UMAP plot representing scRNA-seq data from hatchlings derived from isolated 2-cell blastomeres.

**Supplemental Figure 2.**

(A-B) Schematic of the cleavages of until the 16-cell stage in the wild type embryo and the isolated 4-cell stage macromere, 1H’, immediately after isolation. 1H’ maintains the WT invariant cleavage program in isolation. E50 and E65, 50 and 65 hours post-lay. (C-D) Lineage tracing of 1h’ and 1H’ in the isolated 4-cell stage macromere. Representative images showing the labeling, or lack there of, for major cell types. 2H’ only gives rise to gut, while 2h’ gives rise to epidermis and muscle, as well as neurons and pharynx, the missing cell types (some images duplicated from Figure 2). Photoconverted RFP (magenta) and unconverted GFP (cyan). Scale bar 50 microns.

**Supplemental Figure 3.**

(A) Visualization of chi-square test, comparing proportions of different cell types (p-value < 2.2e-16). Micromere derived embryos have significantly higher proportions of neuronal cell types at 3 dpd. (B-G) Fluorescent *in situ hybridization* for markers for epidermis, gut, neurons, muscle, pharynx, and statocyst in four days post dissection zygote, isolated 2-cell blastomere, isolated 4-cell macromere, isolated 8-cell micromere cap, and isolated 8-cell macromeres. The pharynx is normally derived from 1h^2^ blastomeres but had contribution from 2H blastomeres. The statocyst is marked with an arrow when visible. At least 4 replicates for each sample and gene. Some replicate images from Figure 3E-H. Arrowheads for emphasis of expression. Scale bar 50 microns.

**Supplemental Figure 4.**

(A-D) Representative images for each major cell type group in the reaggregation experiment. (some images duplicated from Figure 4). Labeled cell types (green) and nuclei (blue) with schematics highlighting newly induced cell types (right). The number of replicates is written in the figure.

**Supplemental Table 1.**
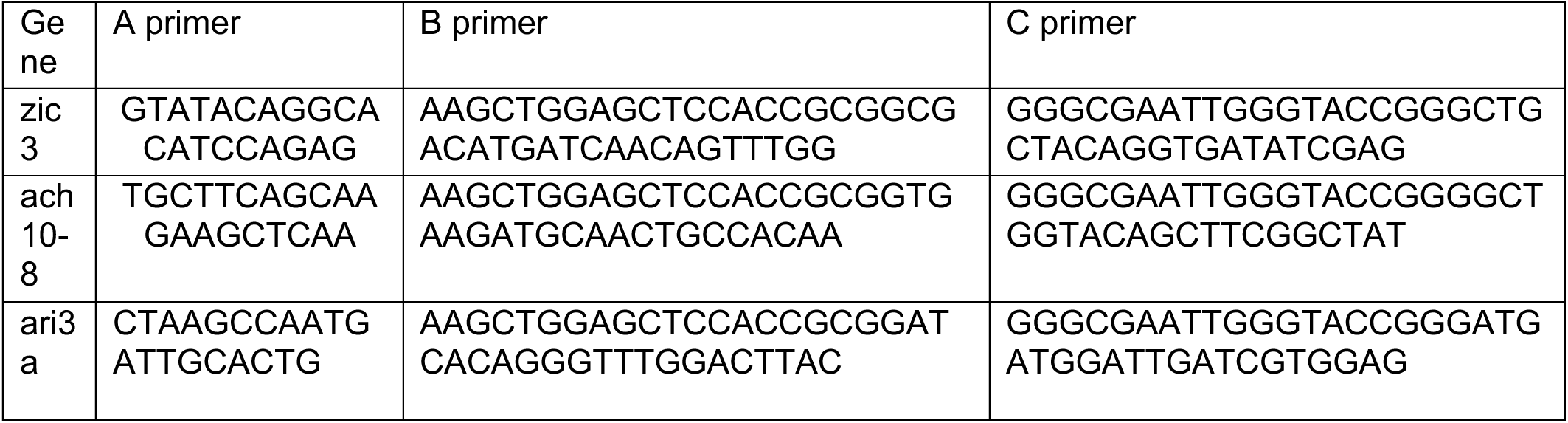
Primer sequences.

## References

Bedzhov, I., Graham, S. J. L., Leung, C. Y., & Zernicka-Goetz, M. (2014). Developmental plasticity, cell fate specification and morphogenesis in the early mouse embryo. Philosophical Transactions of the Royal Society B: Biological Sciences, 369(1657), 20130538. 10.1098/rstb.2013.0538

Benirschke, K. (2009). The monozygotic twinning process, the twin-twin transfusion syndrome and acardiac twins. Placenta, 30(11), 923–928. 10.1016/j.placenta.2009.08.009

Boyer, B. C. (1986). Determinative development in the polyclad turbellarian, Hoploplana inquilina. International Journal of Invertebrate Reproduction and Development, 9(2), 243–251. 10.1080/01688170.1986.10510199

Boyer, B. C., Henry, J. Q., & Martindale, M. Q. (1996). Modified Spiral Cleavage: The Duet Cleavage Pattern and Early Blastomer Fates in the Acoel Turbellarian Neochildia fusca. The Biological Bulletin, 191(2), 285–286. 10.1086/BBLv191n2p285

Bump, P., Loubet-Senear, K., Arnold, S., & Srivastava, M. (2023). Chromatin profiling data indicate regulatory mechanisms for differentiation during development in the acoel Hofstenia miamia (p. 2023.12.05.570175). bioRxiv. 10.1101/2023.12.05.570175

Bushmanova, E., Antipov, D., Lapidus, A., & Prjibelski, A. D. (2019). rnaSPAdes: A de novo transcriptome assembler and its application to RNA-Seq data. GigaScience, 8(9), giz100. 10.1093/gigascience/giz100

Cannon, J. T., Vellutini, B. C., Smith, J., Ronquist, F., Jondelius, U., & Hejnol, A. (2016). Xenacoelomorpha is the sister group to Nephrozoa. Nature, 530(7588), 89–93. 10.1038/nature16520

Conklin, E. G. (1905). Mosaic development in ascidian eggs. Journal of Experimental Zoology, 2(2), 145–223. https://books.google.com/books?hl=en&lr=&id=y7gBAAAAYAAJ&oi=fnd&pg=PA145&dq=mosaic+development+embryo&ots=iHlhJnpDig&sig=E2GVPXpwp7Lcal17K6tozybPRCI

Cunha, T. J., & Giribet, G. (2019). A congruent topology for deep gastropod relationships. Proceedings of the Royal Society B: Biological Sciences, 286(1898), 20182776. 10.1098/rspb.2018.2776

Davidson, E. H. (1990). How embryos work: A comparative view of diverse modes of cell fate specification. Development, 108(3), 365–389. 10.1242/dev.108.3.365

Davidson, E. H. (1991). Spatial mechanisms of gene regulation in metazoan embryos. Development, 113(1), 1–26. 10.1242/dev.113.1.1

Davidson, E. H., Cameron, R. A., & Ransick, A. (1998). Specification of cell fate in the sea urchin embryo: Summary and some proposed mechanisms. Development, 125(17), 3269–3290. 10.1242/dev.125.17.3269

Edgecombe, G. D., Giribet, G., Dunn, C. W., Hejnol, A., Kristensen, R. M., Neves, R. C., Rouse, G. W., Worsaae, K., & Sørensen, M. V. (2011). Higher-level metazoan relationships: Recent progress and remaining questions. Organisms Diversity & Evolution, 11(2), 151–172. 10.1007/s13127-011-0044-4

Egger, B., Steinke, D., Tarui, H., Mulder, K. D., Arendt, D., Borgonie, G., Funayama, N., Gschwentner, R., Hartenstein, V., Hobmayer, B., Hooge, M., Hrouda, M., Ishida, S., Kobayashi, C., Kuales, G., Nishimura, O., Pfister, D., Rieger, R., Salvenmoser, W., … Ladurner, P. (2009). To Be or Not to Be a Flatworm: The Acoel Controversy. PLOS ONE, 4(5), e5502. 10.1371/journal.pone.0005502

Extavour, C. G. (2005). The fate of isolated blastomeres with respect to germ cell formation in the amphipod crustacean Parhyale hawaiensis. Developmental Biology, 277(2), 387–402. 10.1016/j.ydbio.2004.09.030

Gerberding, M., Browne, W. E., & Patel, N. H. (2002). Cell lineage analysis of the amphipod crustacean Parhyale hawaiensis reveals an early restriction of cell fates. Development, 129(24), 5789–5801. 10.1242/dev.00155

Gilbert, D. (2013). Gene-omes built from mRNA-seq not genome DNA.

Grabherr, M. G., Haas, B. J., Yassour, M., Levin, J. Z., Thompson, D. A., Amit, I., Adiconis, X., Fan, L., Raychowdhury, R., Zeng, Q., Chen, Z., Mauceli, E., Hacohen, N., Gnirke, A., Rhind, N., di Palma, F., Birren, B. W., Nusbaum, C., Lindblad-Toh, K., … Regev, A. (2011). Full-length transcriptome assembly from RNA-Seq data without a reference genome. Nature Biotechnology, 29(7), Article 7. 10.1038/nbt.1883

Guralnick, R. P., & Lindberg, D. R. (2001). RECONNECTING CELL AND ANIMAL LINEAGES: WHAT DO CELL LINEAGES TELL US ABOUT THE EVOLUTION AND DEVELOPMENT OF SPIRALIA? Evolution, 55(8), 1501–1519. 10.1111/j.0014-3820.2001.tb00671.x

Haas, B. J., Papanicolaou, A., Yassour, M., Grabherr, M., Blood, P. D., Bowden, J., Couger, M. B., Eccles, D., Li, B., Lieber, M., MacManes, M. D., Ott, M., Orvis, J., Pochet, N., Strozzi, F., Weeks, N., Westerman, R., William, T., Dewey, C. N., … Regev, A. (2013). De novo transcript sequence reconstruction from RNA-seq using the Trinity platform for reference generation and analysis. Nature Protocols, 8(8), Article 8. 10.1038/nprot.2013.084

Hejnol, A., Obst, M., Stamatakis, A., Ott, M., Rouse, G. W., Edgecombe, G. D., Martinez, P., Baguñà, J., Bailly, X., Jondelius, U., Wiens, M., Müller, W. E. G., Seaver, E., Wheeler, W. C., Martindale, M. Q., Giribet, G., & Dunn, C. W. (2009). Assessing the root of bilaterian animals with scalable phylogenomic methods. Proceedings of the Royal Society B: Biological Sciences, 276(1677), 4261–4270. 10.1098/rspb.2009.0896

Hejnol, A., & Schnabel, R. (2005). The eutardigrade Thulinia stephaniae has an indeterminate development and the potential to regulate early blastomere ablations. Development, 132(6), 1349–1361. 10.1242/dev.01701

Henry, J. J., Amemiya, S., Wray, G. A., & Raff, R. A. (1989). Early inductive interactions are involved in restricting cell fates of mesomeres in sea urchin embryos. Developmental Biology, 136(1), 140–153. 10.1016/0012-1606(89)90137-1

Henry, J. Q., Martindale, M. Q., & Boyer, B. C. (2000). The Unique Developmental Program of the Acoel Flatworm, Neochildia fusca. Developmental Biology, 220(2), 285–295. 10.1006/dbio.2000.9628

Holland, L. Z., & Holland, N. D. (2007). A revised fate map for amphioxus and the evolution of axial patterning in chordates. Integrative and Comparative Biology, 47(3), 360–372. 10.1093/icb/icm064

Hörstadius, S. (1957). On the Regulation of Bilateral Symmetry in Plutei with Exchanged Meridional Halves and in Giant Plutei. Development, 5(1), 60–73. 10.1242/dev.5.1.60

Hulett, R. E., Kimura, J. O., Bolaños, D. M., Luo, Y.-J., Rivera-López, C., Ricci, L., & Srivastava, M. (2023). Acoel single-cell atlas reveals expression dynamics and heterogeneity of adult pluripotent stem cells. Nature Communications, 14(1), 2612. 10.1038/s41467-023-38016-4

Hulett, R. E., Potter, D., & Srivastava, M. (2020). Neural architecture and regeneration in the acoel Hofstenia miamia. Proceedings of the Royal Society B: Biological Sciences, 287(1931), 20201198. 10.1098/rspb.2020.1198

Jeffery, W. R. (2001). Determinants of cell and positional fate in ascidian embryos. In International Review of Cytology (Vol. 203, pp. 3–62). Academic Press. 10.1016/S0074-7696(01)03003-0

Kapli, P., Natsidis, P., Leite, D. J., Fursman, M., Jeffrie, N., Rahman, I. A., Philippe, H., Copley, R. R., & Telford, M. J. (2021). Lack of support for Deuterostomia prompts reinterpretation of the first Bilateria. Science Advances, 7(12), eabe2741. 10.1126/sciadv.abe2741

Kapli, P., & Telford, M. J. (2020). Topology-dependent asymmetry in systematic errors affects phylogenetic placement of Ctenophora and Xenacoelomorpha. Science Advances, 6(50), eabc5162. 10.1126/sciadv.abc5162

Katayama, M., Ellersieck, M. R., & Roberts, R. M. (2010). Development of Monozygotic Twin Mouse Embryos from the Time of Blastomere Separation at the Two-Cell Stage to Blastocyst1. Biology of Reproduction, 82(6), 1237–1247. 10.1095/biolreprod.109.082982

Kimura, J. O., Bolaños, D. M., Ricci, L., & Srivastava, M. (2022). Embryonic origins of adult pluripotent stem cells. Cell, 185(25), 4756–4769.e13. 10.1016/j.cell.2022.11.008

Kimura, J. O., Ricci, L., & Srivastava, M. (2021). Embryonic development in the acoel Hofstenia miamia. Development, 148(13), dev188656. 10.1242/dev.188656

Kobayashi, K., Sawada, K., Yamamoto, H., Wada, S., Saiga, H., & Nishida, H. (2003). Maternal macho-1 is an intrinsic factor that makes cell response to the same FGF signal differ between mesenchyme and notochord induction in ascidian embryos. Development, 130(21), 5179–5190. 10.1242/dev.00732

Lambert, J. D. (2010). Developmental Patterns in Spiralian Embryos. Current Biology, 20(2), R72–R77. 10.1016/j.cub.2009.11.041

Laufer, J. S., Bazzicalupo, P., & Wood, W. B. (1980). Segregation of developmental potential in early embryos of caenorhabditis elegans. Cell, 19(3), 569–577. 10.1016/S0092-8674(80)80033-X

Lawrence, P. A., & Levine, M. (2006). Mosaic and regulative development: Two faces of one coin. Current Biology, 16(7), R236–R239. 10.1016/j.cub.2006.03.016

Livingston, B. T., & Wilt, F. H. (1990). Range and stability of cell fate determination in isolated sea urchin blastomeres. Development, 108(3), 403–410. 10.1242/dev.108.3.403

Maduro, M. F. (2010). Cell fate specification in the C. elegans embryo. Developmental Dynamics, 239(5), 1315–1329. 10.1002/dvdy.22233

Marlétaz, F., Peijnenburg, K. T. C. A., Goto, T., Satoh, N., & Rokhsar, D. S. (2019). A New Spiralian Phylogeny Places the Enigmatic Arrow Worms among Gnathiferans. Current Biology, 29(2), 312–318.e3. 10.1016/j.cub.2018.11.042

McCain, E. R., & Cather, J. N. (1989). Regulative and mosaic development of Ilyanassa obsoleta embryos lacking the A and C quadrants. Invertebrate Reproduction & Development, 15(3), 185–192. 10.1080/07924259.1989.9672042

Nishida, H. (2005). Specification of embryonic axis and mosaic development in ascidians. Developmental Dynamics, 233(4), 1177–1193. 10.1002/dvdy.20469

Nishida, H., & Sawada, K. (2001). Macho-1 encodes a localized mRNA in ascidian eggs that specifies muscle fate during embryogenesis. Nature, 409(6821), Article 6821. 10.1038/35055568

Pertea, M., Pertea, G. M., Antonescu, C. M., Chang, T.-C., Mendell, J. T., & Salzberg, S. L. (2015). StringTie enables improved reconstruction of a transcriptome from RNA-seq reads. Nature Biotechnology, 33(3), Article 3. 10.1038/nbt.3122

Philippe, H., Brinkmann, H., Copley, R. R., Moroz, L. L., Nakano, H., Poustka, A. J., Wallberg, A., Peterson, K. J., & Telford, M. J. (2011). Acoelomorph flatworms are deuterostomes related to Xenoturbella. Nature, 470(7333), 255–258. 10.1038/nature09676

Philippe, H., Brinkmann, H., Martinez, P., Riutort, M., & Baguñà, J. (2007). Acoel Flatworms Are Not Platyhelminthes: Evidence from Phylogenomics. PLOS ONE, 2(8), e717. 10.1371/journal.pone.0000717

Price, A. L., Modrell, M. S., Hannibal, R. L., & Patel, N. H. (2010). Mesoderm and ectoderm lineages in the crustacean Parhyale hawaiensis display intra-germ layer compensation. Developmental Biology, 341(1), 256–266. 10.1016/j.ydbio.2009.12.006

Priess, J. R., & Thomson, J. N. (1987). Cellular interactions in early C. elegans embryos. Cell, 48(2), 241–250. 10.1016/0092-8674(87)90427-2

Ricci, L., & Srivastava, M. (2021). Transgenesis in the acoel worm Hofstenia miamia. Developmental Cell, 56(22), 3160–3170. e4.

Rock, A. Q., & Srivastava, M. (2025). The gain and loss of plasticity during development and evolution. Trends in Cell Biology, In press.

Ruiz-Trillo, I., Riutort, M., Fourcade, H. M., Baguñà, J., & Boore, J. L. (2004). Mitochondrial genome data support the basal position of Acoelomorpha and the polyphyly of the Platyhelminthes. Molecular Phylogenetics and Evolution, 33(2), 321–332. 10.1016/j.ympev.2004.06.002

Srivastava, M. (2022). Studying development, regeneration, stem cells, and more in the acoel *Hofstenia miamia*. In B. Goldstein & M. Srivastava (Eds.), Current Topics in Developmental Biology (Vol. 147, pp. 153–172). Academic Press. 10.1016/bs.ctdb.2022.01.003

Srivastava, M., Mazza-Curll, K. L., van Wolfswinkel, J. C., & Reddien, P. W. (2014). Whole-body acoel regeneration is controlled by Wnt and Bmp-Admp signaling. Current Biology, 24(10), 1107–1113.

Sulston, J. E., Schierenberg, E., White, J. G., & Thomson, J. N. (1983). The embryonic cell lineage of the nematode Caenorhabditis elegans. Developmental Biology, 100(1), 64–119. 10.1016/0012-1606(83)90201-4

The UniProt Consortium. (2023). UniProt: The Universal Protein Knowledgebase in 2023. Nucleic Acids Research, 51(D1), D523–D531. 10.1093/nar/gkac1052

Wilt, F. H. (1987). Determination and morphogenesis in the sea urchin embryo. Development, 100(4), 559–576. 10.1242/dev.100.4.559

Yamaguchi, M., Kinoshita, T., & Ohba, Y. (1994). Fractionation of Micromeres, Mesomeres, and Macromeres of 16-cell Stage Sea Urchin Embryos by Elutriation*. Development, Growth & Differentiation, 36(4), 381–387. 10.1111/j.1440-169X.1994.00381.x

Zilionis, R., Nainys, J., Veres, A., Savova, V., Zemmour, D., Klein, A. M., & Mazutis, L. (2017). Single-cell barcoding and sequencing using droplet microfluidics. Nature Protocols, 12(1), 44–73. 10.1038/nprot.2016.154

