## Supplementary figures and images for "Totipotency and high plasticity in an embryo with a stereotyped, invariant cleavage program"

### Supplemental Figures

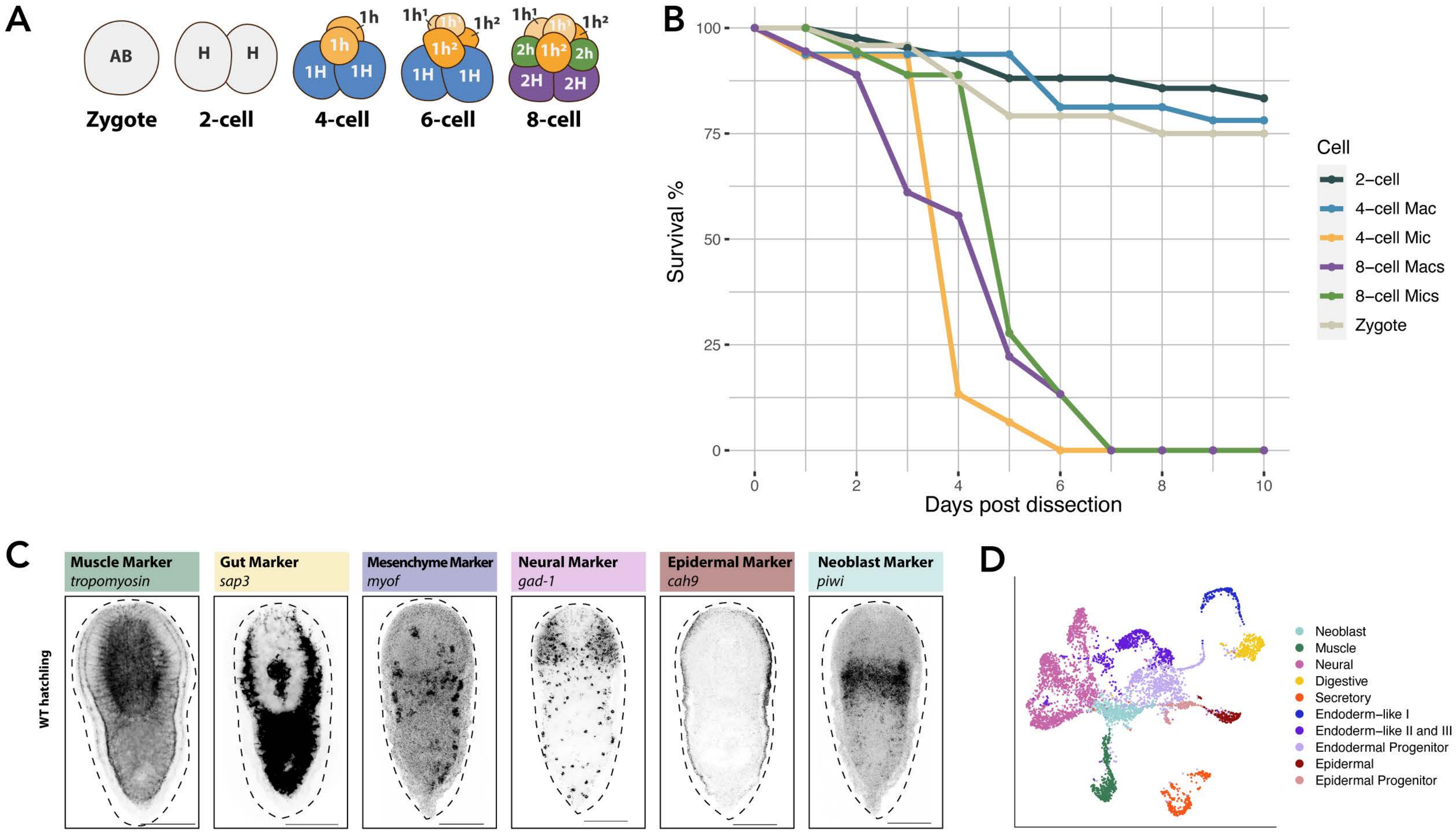


Supplemental Figure 1

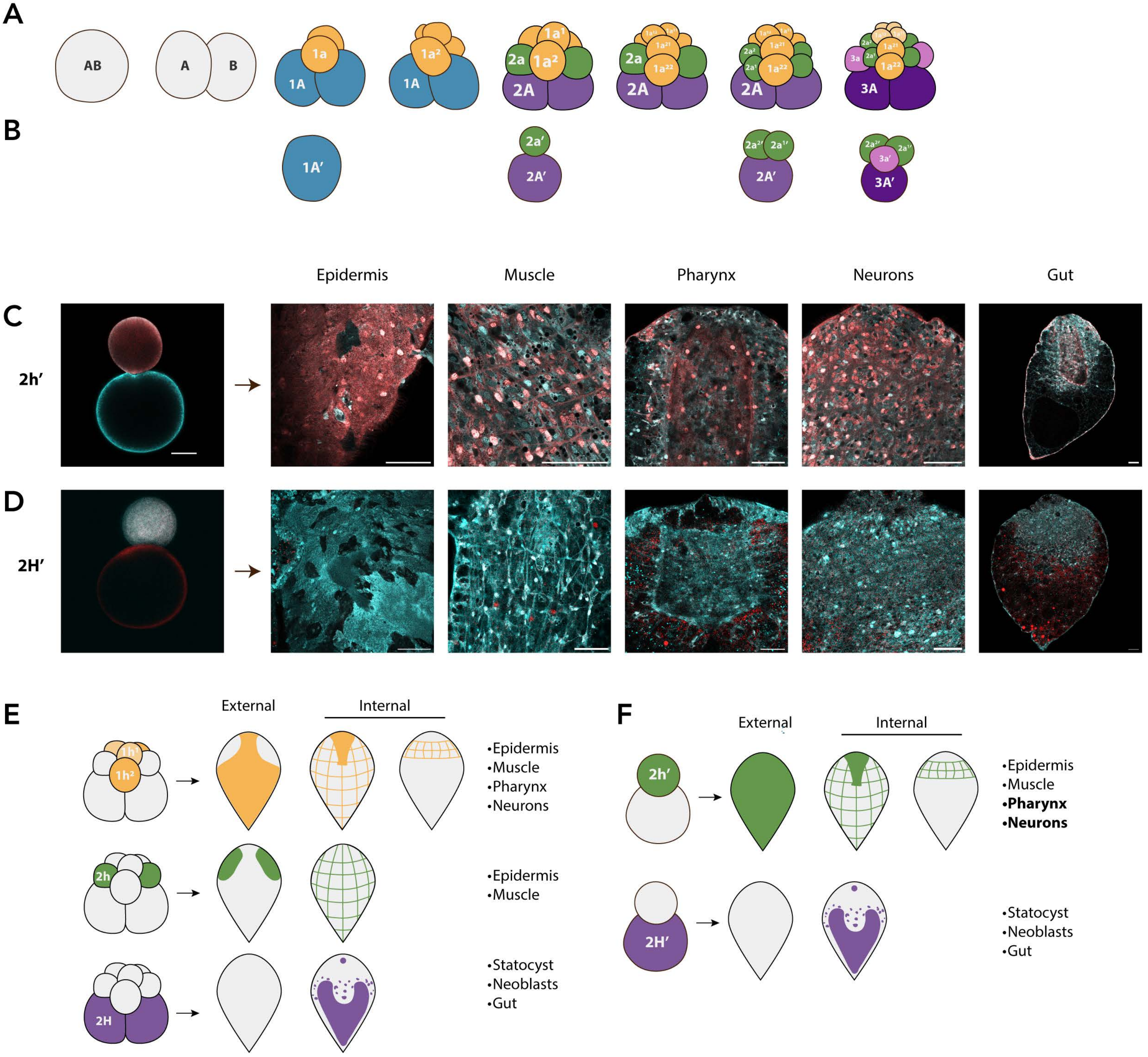


Supplemental Figure 2

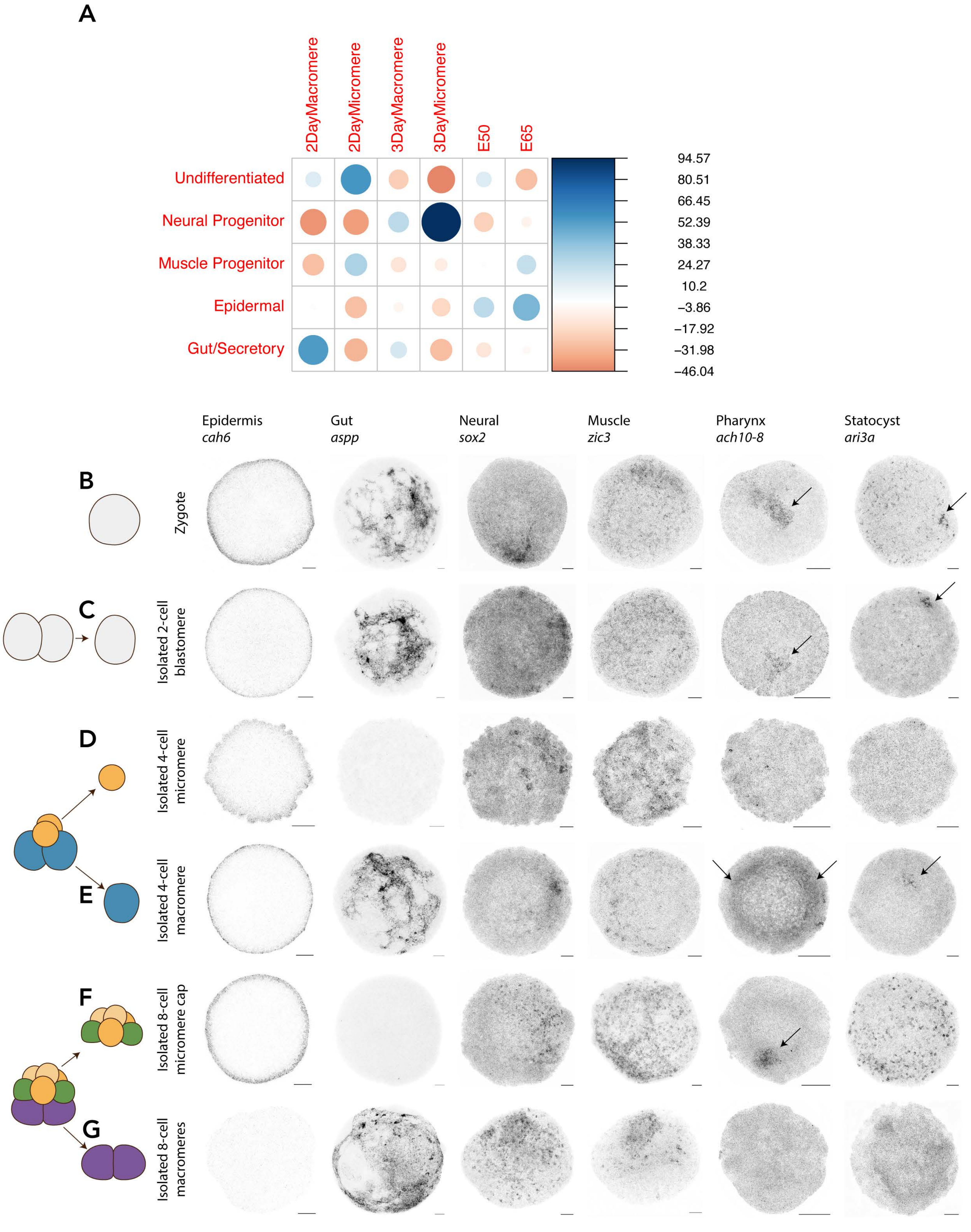


Supplemental Figure 3

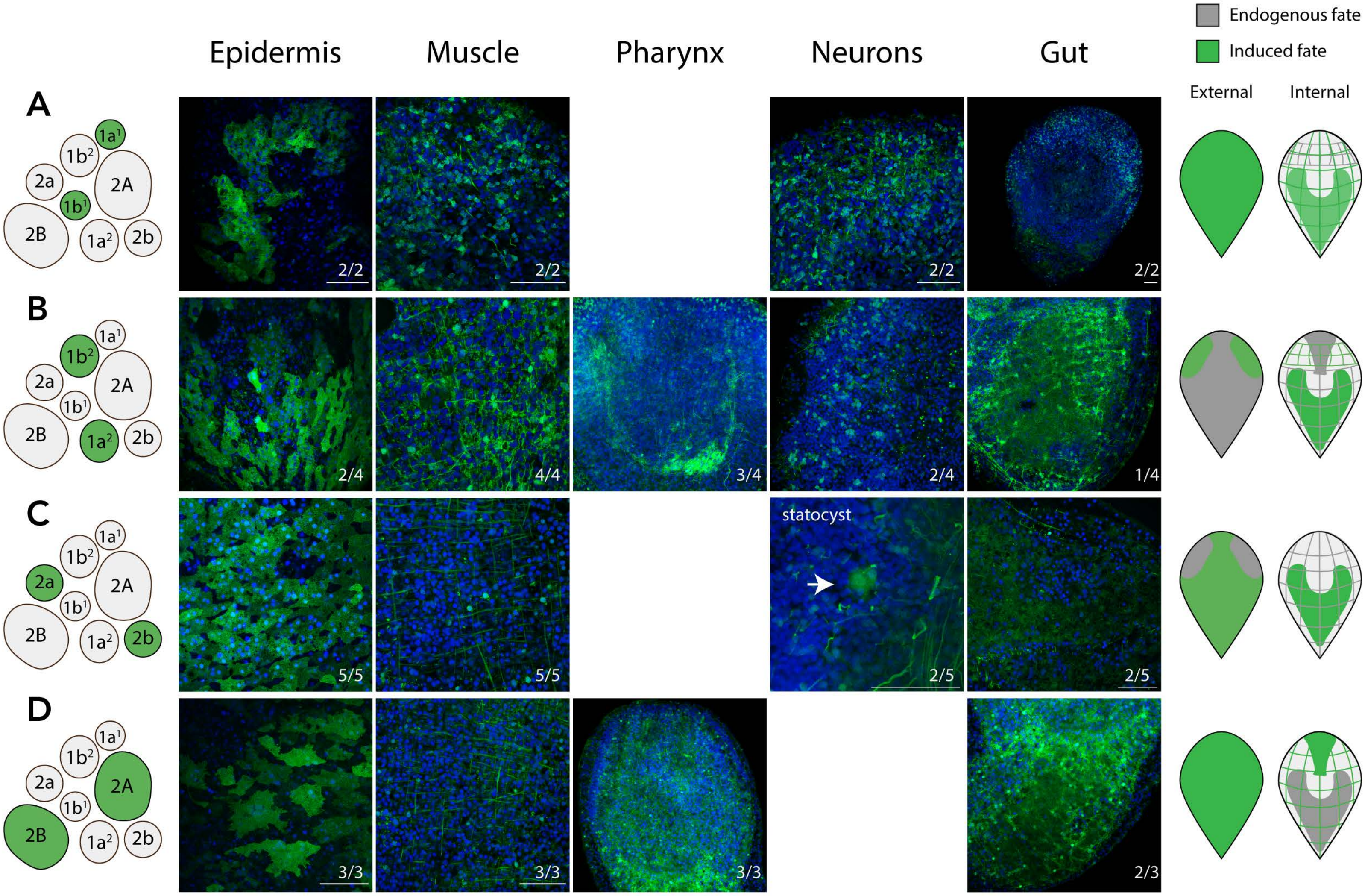


**Supplemental Figure 4**
